# Native dynamics and allosteric responses in PTP1B probed by high-resolution HDX-MS

**DOI:** 10.1101/2023.07.12.548582

**Authors:** Virgil A. Woods, Rinat R. Abzalimov, Daniel A. Keedy

## Abstract

Protein tyrosine phosphatase 1B (PTP1B) is a validated therapeutic target for obesity, diabetes, and certain types of cancer. In particular, allosteric inhibitors hold potential for therapeutic use, but an incomplete understanding of conformational dynamics and allostery in this protein has hindered their development. Here, we interrogate solution dynamics and allosteric responses in PTP1B using high-resolution hydrogen-deuterium exchange mass spectrometry (HDX-MS), an emerging and powerful biophysical technique. Using HDX-MS, we obtain a detailed map of the solution dynamics of apo PTP1B, revealing several flexible loops interspersed among more constrained and rigid regions within the protein structure, as well as local regions that exchange faster than expected from their secondary structure and buriedness. We demonstrate that our HDX rate data obtained in solution adds value to predictions of dynamics derived from a pseudo-ensemble constructed from ∼200 crystal structures of PTP1B. Furthermore, we report HDX-MS maps for PTP1B with active-site vs. allosteric small-molecule inhibitors. These maps reveal distinct, dramatic, and widespread effects on protein dynamics relative to the apo form, including changes to dynamics in locations distal (>35 Å) from the respective ligand binding sites. These results help shed light on the allosteric nature of PTP1B and the surprisingly far-reaching consequences of inhibitor binding in this important protein. Overall, our work showcases the potential of HDX-MS for elucidating protein conformational dynamics and allosteric effects of small-molecule ligands, and highlights the potential of integrating HDX-MS alongside other complementary methods to guide the development of new therapeutics.

## Introduction

The protein tyrosine phosphatase (PTP) family is a diverse group of enzymes that regulate cellular signaling pathways by catalyzing the dephosphorylation of phosphotyrosine (pTyr) residues on proteins. The human PTP family contains more than 100 members that can be classified into four subfamilies based on their domain organization and function ^1^. PTP1B is the archetypal member of the PTP family, characterized by its conserved catalytic domain, as well as a non-conserved, disordered C-terminal domain that regulates its intracellular localization and activity. PTP1B has been extensively studied due to its central role in regulating insulin and leptin signaling and its potential as a therapeutic target for metabolic disorders. Structural and dynamic studies of PTP1B have provided insights into the mechanisms underlying its catalytic activity, substrate recognition, and regulation. These insights have been used to guide the design of various small-molecule inhibitors targeting PTP1B. In addition, comparative studies of PTP1B and other members of the PTP family have revealed conserved structural and dynamical features that can be exploited in drug discovery efforts targeting other PTP family members ^2^.

The catalytic mechanism of PTP1B involves a conserved cysteine residue, Cys215, within the active-site P loop, which acts as a nucleophile and forms a covalent thiol-phosphate intermediate with the target protein’s pTyr residue ^3^. The catalytic activity of PTP1B is also facilitated by several other critical loops in the protein’s structure, particularly the WPD loop and Q loop. The WPD loop, which includes a highly conserved aspartic acid residue, Asp181, acts as an acid and protonates the dephosphorylated tyrosine group, leading to its release ^4^. Arg221, located spatially adjacent to the WPD loop at the end of the P loop, also forms hydrogen bonds with the target protein’s pTyr residue, maintaining PTP1B in a catalytically active conformation ^5^. The Q loop, which includes a highly conserved glutamine residue, Gln262, activates a water molecule that attacks the phosphocysteine intermediate, releasing inorganic phosphate and regenerating the catalytic cysteine ^6,7^. These critical loops are essential for the efficient catalytic activity of PTP1B and are located in protein regions subject to conformational changes upon binding of inhibitors.

Despite its therapeutic potential, drugging the active site of phosphatases like PTP1B has proven challenging due to these enzymes’ highly polar and conserved nature. This negatively affects the ability of small molecules to exert specific activity against particular phosphatases and achieve drug-like membrane permeability needed to reach these largely cytoplasmic enzymatic domains. TCS401 is an active-site inhibitor of PTP1B that has been extensively used in structural studies ^8,9^. These studies have revealed the critical interactions between TCS401 and the active site of PTP1B, providing essential insights into the design of active-site inhibitors. Alternatively, allosteric inhibitors boast potential advantages in both bioavailability and phosphatase specificity by targeting less polar and less conserved binding sites. BB3 is an allosteric inhibitor of PTP1B that binds to a distal site known as the BB site ^10^. A few isolated locations within PTP1B appear to undergo conformational changes in structural studies of apo versus liganded conditions with these two distinct inhibitors: TCS401 binding closes the critical WPD loop, which is observed to cause ordering of the key allosteric α7 helix, while BB3 binds in proximity to the α7 helix and thereby displaces its ordered conformation, promoting the open state of the WPD loop (**Fig. 1**). Although these studies have yielded valuable insights into mechanisms of PTP allostery and inhibition, a deeper understanding of the conformational changes and protein dynamics that underlie these processes is necessary to design more effective inhibitors that can selectively target PTP1B without affecting other members of the PTP family.

**Figure 1:**
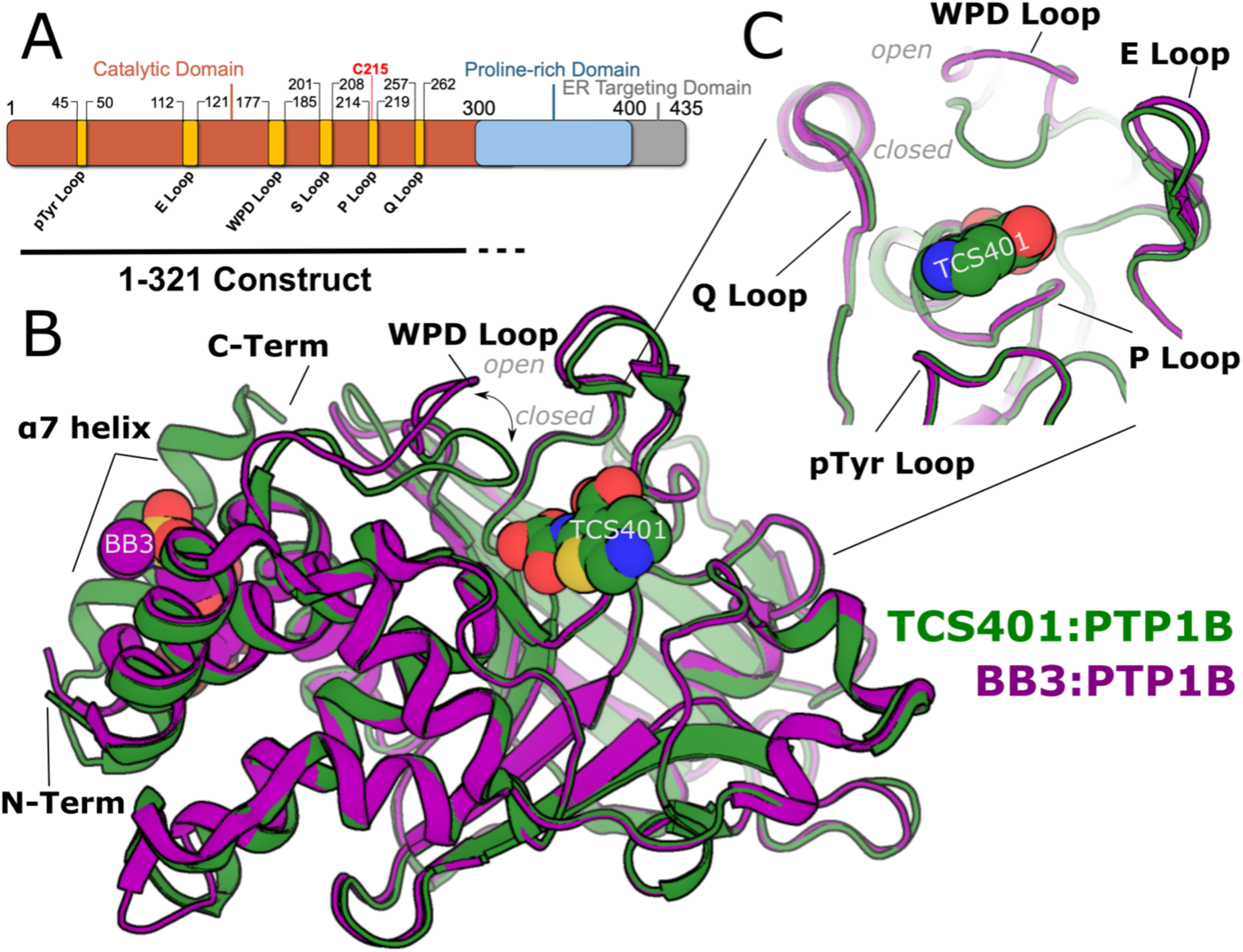
Overview of PTP1B structure and key sites. **A.** Schematic of full-length PTP1B primary structure, including catalytic domain and disordered C-terminus. Key structural sites in the catalytic domain are labeled. The 1-321 construct used throughout this study is indicated below the diagram. The last ∼22 residues of this construct do not appear in crystal structures. **B.** Superposed structures of the PTP1B catalytic domain bound to the active-site inhibitor TCS401 (green, PDB ID 5K9W, closed state) and bound to the allosteric inhibitor BB3 (purple, PDB ID 1T49, open state). **C.** Close-up view of PTP1B active site showing the positions of catalytic loops surrounding TCS401.

Previous efforts to measure PTP1B conformational dynamics have included a variety of biophysical methods. Crystal structures of PTP1B in complex with allosteric inhibitors have revealed that binding at an allosteric site can modulate the conformation of the active site, leading to enzymatic inhibition ^9,11^. However, the structural changes observed by crystallography may only partially capture the dynamic nature of the protein in solution. Nuclear magnetic resonance (NMR) spectroscopy has also provided insights into structural regions with motions at similar timescales as well as remote responses to ligands and mutations ^9,12–14^. Complementary to these established methods, hydrogen-deuterium exchange mass spectrometry (HDX-MS) measures the exchange of labile hydrogen atoms on a protein with deuterium in solution, which is then detected by mass spectrometry. Previously, PTP1B dynamics in solution were studied using only low-resolution HDX-MS ^15^. However, as a result of technological advances, high-resolution local HDX-MS is emerging as a powerful tool for obtaining unique insights into protein dynamics and conformational changes ^16^. This more mature technique provides detailed information about a protein’s local flexibility and stability, which can reveal conformational changes induced by ligand binding or protein-protein interactions. Importantly, HDX-MS can probe protein regions that may be more difficult to characterize by other methods, such as disordered regions or flexible loops.

In this study, we use high-resolution local HDX-MS to explore the dynamics of PTP1B in the apo form vs. upon binding to the active-site inhibitor TCS401 or the allosteric inhibitor BB3. Our results provide new insights into the mechanisms of inhibition by these compounds, and highlight the utility of HDX-MS for studying protein dynamics and allosteric regulation, with potential implications for the development of effective PTP1B inhibitors.

## Results

### Overview of PTP1B structure

The overall structure of the catalytic domain of PTP1B consists of several crucial catalytic loops, designated the WPD, Q, pTyr, E, Q, and P loops (**Fig. 1**). In addition to these loops, a distal α helix (α7), located approximately 20 Å away from the catalytic Cys215, has been implicated as a key element in the allosteric regulation of PTP1B ^9,11,17^. Notably, the conformation of the α7 helix is influenced by the state of the WPD loop: when the WPD loop is closed, α7 becomes ordered; conversely, when the WPD loop is open, α7 becomes disordered.

Previous studies have shown that active-site inhibitors engage nearly all of the catalytic loops directly, including the binding of TCS401 within the pocket formed by the WPD, pTyr, E, Q, and P loops. This specific interaction between TCS401 and the catalytic loops parallels their critical role in PTP1B’s enzymatic function. In contrast, some allosteric ligands, such as BB3, induce an allosteric binding event that displaces the α7 helix, stabilizing it in a disordered state. This binding event has been observed to specifically stabilize the open conformation of the catalytic WPD loop (**Fig. 1B**). Notably, while the crystal structure of PTP1B bound to compound BB3 reveals the conformational change in the WPD loop and residues along an allosteric pathway linking these two sites, it indicates little to no change in the conformation of active-site residues, nor indeed of any other residues beyond the proposed allosteric pathway ^10^.

### HDX-MS of apo PTP1B

To investigate the solution dynamics of the catalytic domain of PTP1B, we performed high-resolution local HDX-MS of the apo protein, including correction for back exchange. As with all experiments in this study, we used a construct of PTP1B containing residues 1-321 (**Fig. 1A**), which is a commonly used truncated construct that constitutes the largest previously used in X-ray crystallography. This experiment represents the first HDX-MS peptide mapping of PTP1B with such high resolution, with over 200 overlapping peptides in the final high-quality peptide map, and 100% sequence coverage, excluding proline residues (which lack a backbone amide group) and the first two residues of the protein sequence (**Fig. 2A**).

**Figure 2:**
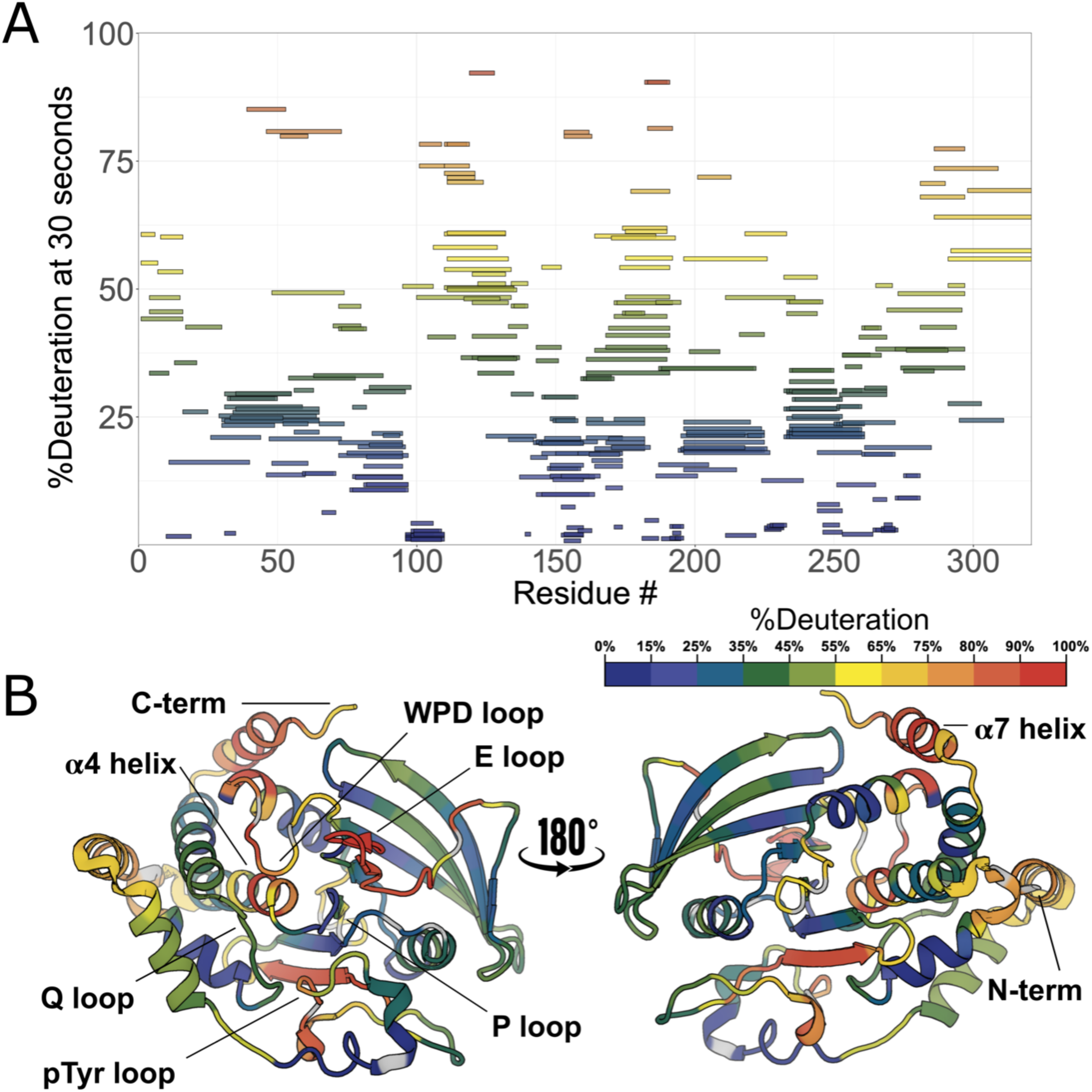
High-resolution local HDX-MS map for apo PTP1B. **A.** Woods plot of HDX rates for 312 peptides of apo PTP1B showing %deuteration at 30 seconds of labeling for each peptide. **B.** Deconvoluted HDX rates at 30 s of labeling time mapped to a crystal structure of the PTP1B catalytic domain in the closed state (PDB ID 1SUG). Several key loops and the α4 helix (residues 187-200) and α7 helix (residues 285-298) are indicated.

The overall HDX-MS profile of PTP1B shows that, under these conditions, many portions of PTP1B are very stable, interrupted by at least eight regions that can best be described as dynamic loops (**Fig. 2A-B**). The regions of low dynamics in the apo state are primarily confined to rigid structural elements such as α helices and β sheets. Peptides corresponding to these loci were shown to substantially deuterate only after incubating for 8 days at 25°C (fully deuterated condition; see Methods). The peptides containing the active-site catalytic cysteine (Cys215) show among the highest degree of stability, with less than 20% of theoretical maximum deuteration happening at the longest time point of 10,000 seconds (2 hours and 47 minutes). In contrast, the active-site WPD loop shows some of the highest dynamics ^12^, reaching above 50% deuteration at just 30 seconds.

The disordered C-terminal region ^13^ begins around residue 299 and continues to the end of the sequence in our construct. As expected, in our local HDX data, this region, which has a high density of proline residues, exhibits a very high degree of dynamics compared to the ordered catalytic domain (**Fig. 2A-B**). As noted above, the α7 helix (residues 285-298), the last quasi-ordered region of the catalytic domain before the disordered tail, has been previously shown to exhibit conditional disorder: it is ordered when the WPD loop is closed, and disordered when the WPD loop is open ^9,11^. Indeed, α7 is quite dynamic in our HDX data. Surprisingly, the residues immediately surrounding Trp291 within α7 are particularly dynamic, even more so than upstream or downstream in the rest of the helix. This observation may be explained by the facts that the Trp291 side chain is a dynamic α7 anchor of sorts that can be displaced via molecular mimicry by the allosteric inhibitor BB3 ^10^ and that this region of α7 is capable of reordering into non-helical conformations in contact with the catalytic domain ^11^.

Two other regions have somewhat higher exchange rates than expected in apo PTP1B. First, the N-terminal region of the α4 helix (residues 221-226), immediately following the catalytic P loop, is buried in the structure of PTP1B. However, in our HDX profile, this region exchanges mildly faster than one might expect, despite being flanked by relatively rigid regions (**Fig. 2A-B**). This observation may be related to the observation that several mutants in this area, including F225Y, increase enzyme activity by modulating latent conformational dynamics ^14^. Second, one of the two edge strands of the main β sheet (residues 69-74) exchanges faster than one might expect for secondary structure, perhaps due to its exposure to solvent.

### Conformational flexibility from crystal structure pseudo-ensemble

In an attempt to attribute structural details to the observed HDX measurements for apo PTP1B, we compared the %deuteration at the earliest time point (30 s) to measures of flexibility calculated from crystal structures, taking advantage of the wealth of available structures for this protein. We analyzed 199 structures of PTP1B from the Protein Data Bank ^18^ (see Methods) in a variety of different experimental conditions, many with different small-molecule ligands, and spanning several different crystal space groups. These diverse structures collectively represent a pseudo-ensemble (**Fig. 3A**) that we hypothesized may represent an estimate of the accessible conformational landscape for PTP1B ^19,20^.

**Figure 3:**
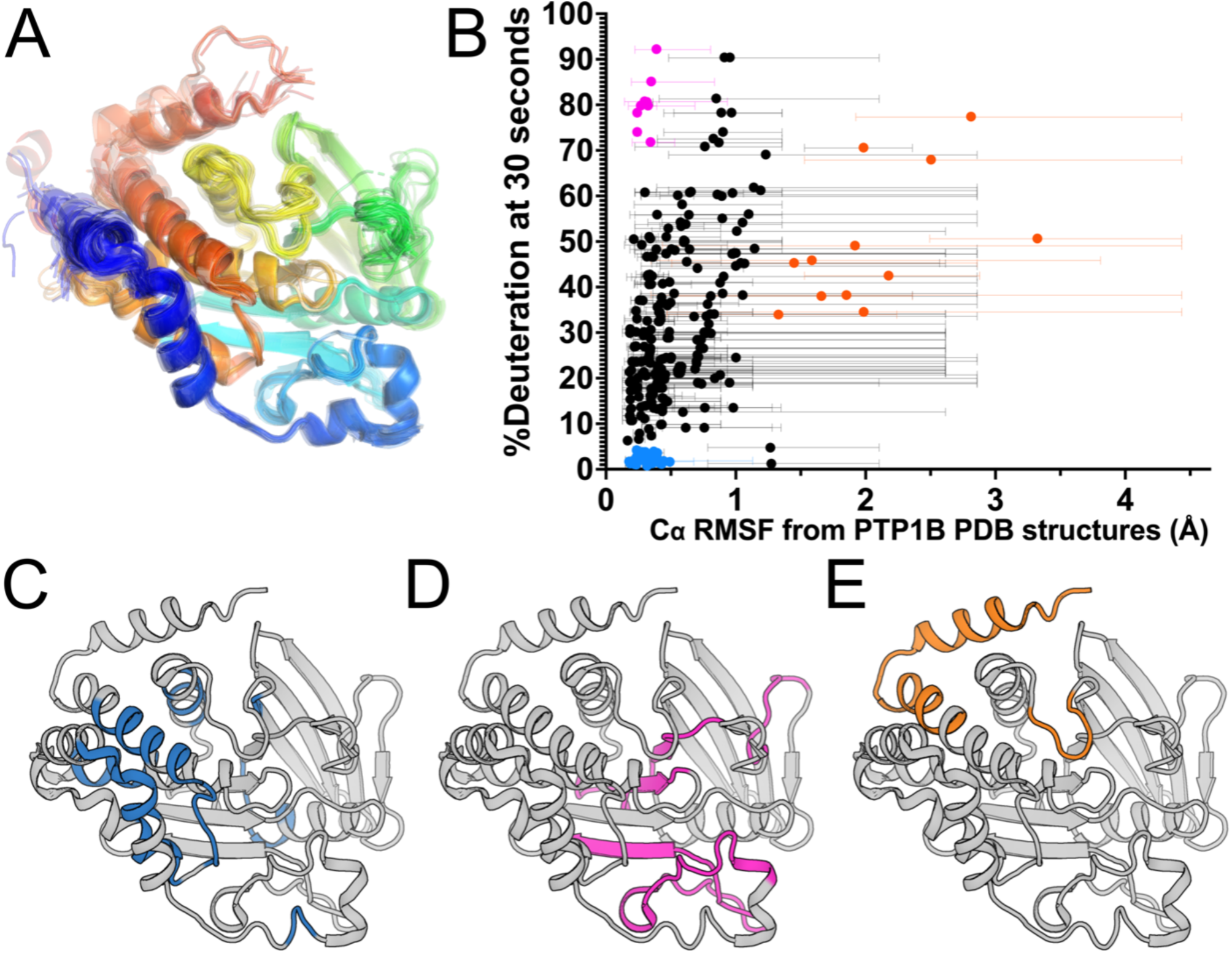
Apo PTP1B local HDX-MS reaction rates are only partially explained by a pseudo-ensemble of crystal structures. **A.** PTP1B pseudo-ensemble derived from all non-PanDDA ^21^ crystal structures from the PDB (n=199; see Methods). Only protein chain A shown; colored from N- to C-terminus (blue to red). **B.** Plot of peptide-level HDX-MS %deuteration (back-exchange corrected) at the 30 second time point vs. pseudo-ensemble Cα root-mean-square fluctuation (RMSF). Average RMSF values are shown as circles. Range of individual residue RMSF values within each peptide are shown as bars **C-E.** The three peptide groups based on criteria of Cα RMSF and HDX from panel **B**, mapped to a crystal structure of the PTP1B catalytic domain in the closed state (PDB ID 1SUG). **C.** Peptides with low mean RMSF (< 0.5 Å) and low HDX (< 5%), in blue. **D.** Peptides with low mean RMSF (< 0.5 Å) and high HDX (> 70%), in magenta. **E.** Peptides with high mean RMSF (> 1.25 Å), in orange.

Using this pseudo-ensemble, we calculated Cα root-mean-square fluctuation (RMSF) for each residue, and compared these RMSF values to local HDX values for apo PTP1B (**Fig. 3B**). Among peptides with high mean Cα RMSF, all have high HDX values. Most of these peptides are from sites previously noted to exhibit conformational heterogeneity in crystal structures, such as the quasi-disordered C-terminal α7 helix and bistable WPD loop (**Fig. 3E**, orange points in **Fig. 3B**). However, among peptides with lower mean Cα RMSF, there is a wide spread of HDX values, following a general pattern of positive correlation, with the exception of two distinct groups. Some peptides with low mean RMSF in the pseudo-ensemble had some of the highest rates of exchange (**Fig. 3D**, magenta points in **Fig. 3B**). These peptides are largely centered on the β strands and loops surrounding the catalytic site. Among these is a peptide (residues 48-73) that includes part of the surprisingly dynamic β strand identified in the apo map of PTP1B (**Fig. 2**). By contrast, peptides with especially low RMSF and exchange were largely centered around the α-helical cluster (**Fig. 3C**, blue points in **Fig. 3B**). Together, these results suggest that more pronounced solution dynamics are qualitatively well predicted by a crystal structure pseudo-ensemble — but for residues that appear similar in such a pseudo-ensemble, high-resolution HDX data are useful to quantitatively differentiate their dynamics.

### HDX-MS with two distinct inhibitors

Having characterized local HDX of apo PTP1B, we next wished to examine changes in local HDX at high spatial resolution induced by distinct small-molecule inhibitors targeting the active site vs. an allosteric site. To do so, we collected local HDX datasets in the presence of the small-molecule inhibitors TCS401 and BB3 (**Fig. S1**). The exchange time points and protocol for protein digestion were identical to those for the apo protein. By comparing the local exchange rate of each amino acid in the presence vs. absence of each inhibitor, we aimed to elucidate the specific effects of each inhibitor on PTP1B dynamics.

Difference HDX Woods plots were produced by coloring peptides based on the difference (change in %deuteration) between the protein-ligand complex and the apo state deuteration levels at every time point. The results show dramatic differences, both upon binding of each ligand and between the different ligands (**Fig. 4A-B**). The subsequent sections of this study delve into a detailed exploration of these observed differences, shedding light on the specific regions that experience changes in dynamics and/or conformation upon binding of each inhibitor.

**Figure 4:**
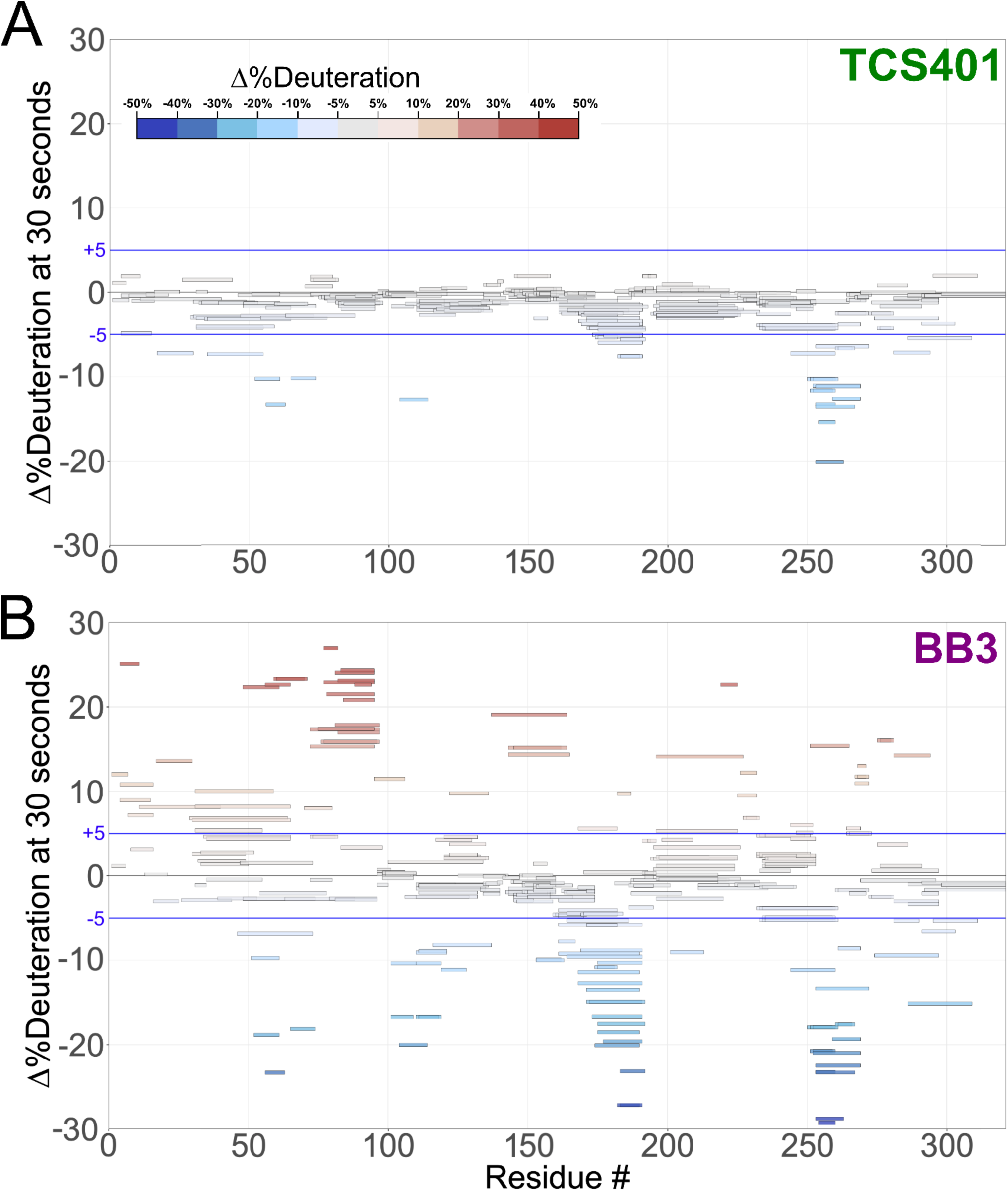
Difference Woods plot of active-site and allosteric inhibitors relative to apo PTP1B. Difference in %deuteration values at 30 seconds in each inhibitor-bound state minus the apo PTP1B state, plotted against amino acid sequence. Blue lines indicate a difference interval of ± 5%. **A.** Active-site inhibitor, TCS401. **B.** Allosteric inhibitor, BB3.

### HDX-MS with active-site inhibitor (TCS401)

As an active-site inhibitor, TCS401 does not rely on changes to the conformation or dynamics of protein residues distal to the active site to convey inhibition. Consistent with this expectation, most of the residues affected by TCS401 binding in the local HDX difference map (**Fig. 4A**, **Fig. S1**) are in active-site loops. These residues exhibit decreased exchange upon inhibitor binding, which may be due to steric blockage of exchanging deuterons by the ligand itself and/or rigidification of the active-site environment upon binding.

The C-terminus of the α5 helix (residues 253-263) exhibits a significant decrease in dynamics (**Fig. 5A**). The effect on dynamics in this region is surprising because α5 is not part of the active site, and previous crystal structures of apo vs. TCS401-bound PTP1B were similar in this area. However, in the amino acid sequence, α5 immediately precedes the active-site Q loop, which also exhibits a decrease in dynamics; thus this local cluster within PTP1B may exhibit correlated dynamics.

**Figure 5:**
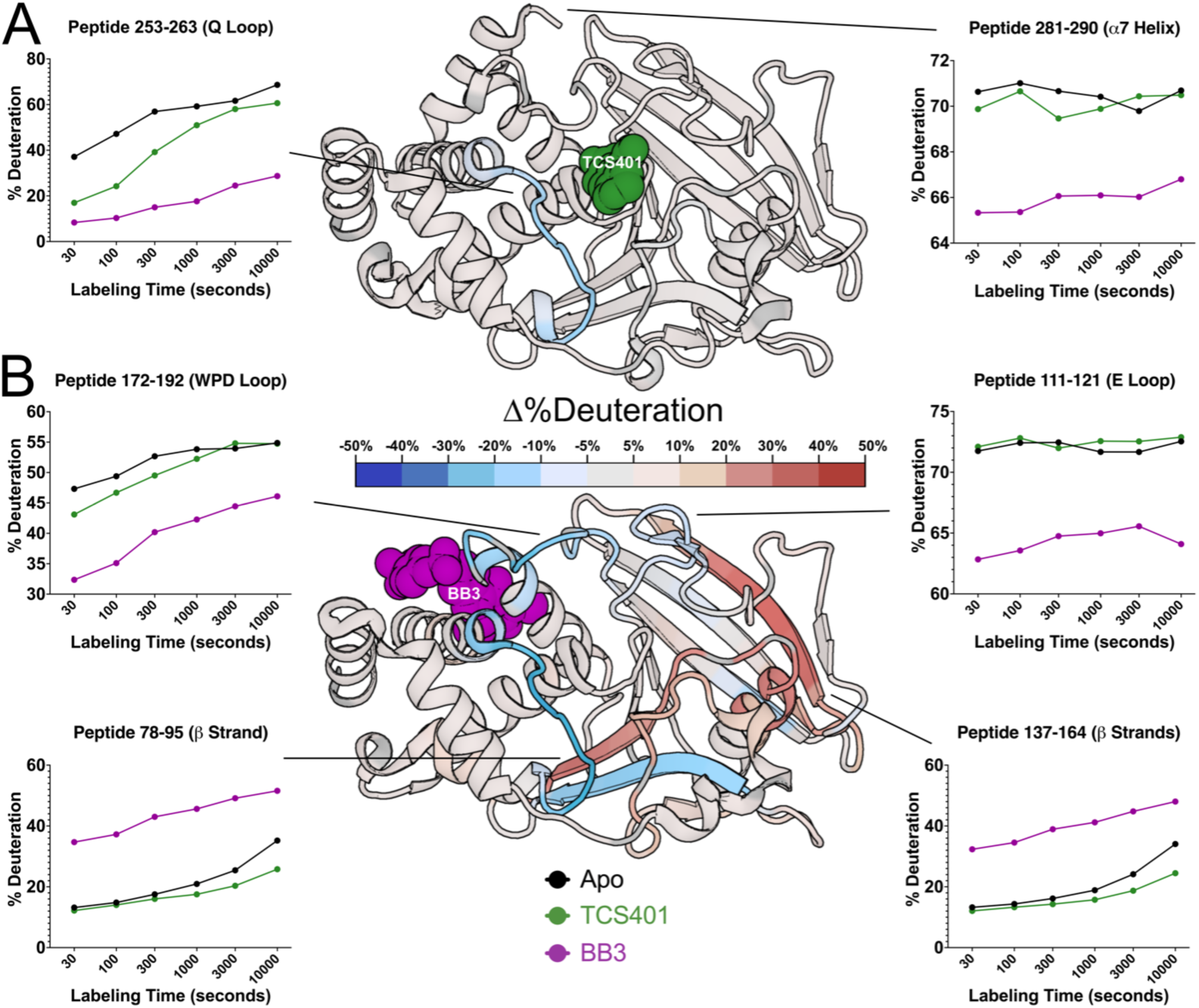
Effects of active-site inhibitor (TCS401) and allosteric site inhibitor (BB3) on local HDX. The structure of PTP1B in complex with **(A)** the active-site inhibitor TCS401 (PDB ID 5K9W) and **(B)** the allosteric inhibitor BB3 (PDB ID 1T49) color-mapped with the deconvoluted differential HDX values at 30 seconds of labeling time. Particular peptides of interest have been selected and their fractional deuterium build-up plots shown. See also **Fig. S3**.

Interestingly, this region also approximately flanks the known secondary pTyr binding site of PTP1B ^22^ (**Fig. S2**). One potential explanation for this observation is that the competitive inhibitor TCS401 binds to both pTyr binding sites, albeit with different affinities. Another possibility is that binding of pTyr (or a pTyr competitor like TCS401) allosterically modulates conformational dynamics at and near the second pTyr site.

### HDX-MS with allosteric inhibitor (BB3)

To determine whether PTP1B responds similarly vs. distinctly to inhibitors at different sites that operate by different mechanisms, we used the allosteric inhibitor BB3 (**Fig. 4B**, **Fig. S1**). As with TCS401, BB3 binding appears to confer the largest protection in the region of the active-site Q loop and flanking α helices (residues 253-263) (**Fig. 4B**). However, compared to TCS401, the effects of BB3 are much more extensive, spanning more sites in the protein, and with a greater variety of effects (**Fig. 4A-B**, **Fig. S1**).

Perhaps most dramatically, unlike with TCS401, several regions have significantly increased exchange upon BB3 binding. This includes residues 79-101 (central β strand and α2 helix) and 139-149 (edge β strand) (**Fig. 4B**, **Fig. S1**). Interestingly, although they are dispersed throughout the primary structure, these regions are all essentially contiguous in the tertiary structure (**Fig. 5B**). The increases in protein flexibility at these sites, which are quite distal from BB3 itself (up to 35 Å), evoke recently reported increases in protein conformational entropy at distal sites upon ligand binding ^23^. Notably, within the one large β sheet in PTP1B, the aforementioned two strands with increased exchange are located amidst several other neighboring strands with more modest changes in exchange (**Fig. 5B**), suggesting that this β sheet undergoes complex dynamical responses ^24^. Unlike at the 30 second time point, the TCS401-bound PTP1B condition at 10,000 seconds shows significant protection of the HDX reaction at the same β strand and α2 helix, which appear more dynamic in BB3-bound PTP1B at all time points (**Fig. S3**).

Unlike the regions described above, several other regions in PTP1B have decreased exchange upon BB3 binding. First, the region with the biggest decrease in exchange upon BB3 binding is the active-site Q loop (**Fig. 4B**, **Fig. 5B**). Interestingly, this region also had the biggest decrease in exchange upon TCS401 binding, marking the greatest similarity between the structural responses to the two ligands (**Fig. 4A-B**, **Fig. 5A-B**). However, with BB3, the regions flanking the Q loop (in α5 and α6) have modest compensatory increased exchange (**Fig. 4B**, **Fig. S1**). Notably, the distal end (C-terminus) of α6 contacts BB3 directly, but it has a substantial decrease in exchange; this suggests a complex pattern of short/medium/long-range allosteric responses to ligand binding.

Second, the WPD loop (residues 178-188) has substantially decreased exchange upon BB3 binding (**Fig. 4B**, **Fig. 5B**, **Fig. S1**). This is perhaps counterintuitive as BB3 is thought to lock the WPD loop in the open state, which one might anticipate would have increased exchange due to higher solvent exposure. However, the observed decrease in exchange may be due to a decrease in open/closed loop dynamics ^12^, supporting the idea that HDX measurements serve as useful proxies for conformational dynamics.

Third, related to the WPD loop, the α7 helix does not have increased exchange, and indeed has mildly decreased exchange, upon BB3 binding (**Fig. 5B**). This is also perhaps counterintuitive as opening of the WPD loop is thought to destabilize the ordered helical state of α7, leading to a disordered ensemble ^11^ that would presumably exchange more quickly. However, our data here are consistent with the idea that the “disordered” α7 may re-engage with the catalytic domain atop the BB site in a quasi-ordered fashion, as supported by previous observations from NMR and room-temperature crystallography ^11,13^.

## Discussion

This study provides detailed maps revealing evidence of conformational dynamics for the archetypal tyrosine phosphatase, PTP1B, using high-resolution local hydrogen-deuterium exchange mass spectrometry (HDX-MS). For the apo protein (**Fig. 2**), many of the patterns are as expected — e.g. most surface-exposed loops undergo greater exchange than most buried regions — but the higher spatial resolution and multiple time points of our new data provide a window into these relative dynamics in unprecedented detail. Other observations from the apo HDX map are more surprising. For example, we observe surprisingly high exchange for a β strand, as well as unexpected differences in the exchange of different portions of the α7 helix, adding nuance to our understanding of this key allosteric hub in PTP1B ^9,11,17^.

In addition to the apo map, our liganded vs. apo difference maps reveal areas of PTP1B that are susceptible to local and distal effects of inhibitor binding (**Fig. 4**, **Fig. 5**). First, upon binding of the active-site inhibitor TCS401, we observed predominantly decreased deuterium uptake, particularly in the region surrounding the Q loop’s interaction with the ligand (**Fig. 4A**, **Fig. 5A**). Interestingly, this region is adjacent to the secondary pTyr binding site in PTP1B, suggesting possible functional relevance to these changes in dynamics (**Fig. S2**).

In contrast to TCS401, binding of the allosteric inhibitor BB3 induced increased hydrogen-deuterium exchange in many parts of the protein (**Fig. 4B**, **Fig. 5B**). Distal from the binding site, several regions that are dispersed in the primary structure (**Fig. 4B**) but nearby in the tertiary structure (**Fig. 5B**) exhibited increased exchange upon ligand binding. This includes residues as far as 35 Å from BB3 itself that were not previously implicated in the BB3 allosteric mechanism ^10^, thus illustrating the power of HDX-MS to uncover surprising aspects of protein conformational dynamics and allostery. In addition to these regions with increased exchange, the Q loop region exhibited decreased exchange upon binding of BB3, even more pronounced than the response to TCS401. These observations suggest that, despite their completely different mechanisms, the two ligands studied here may modulate some similar aspects of the PTP1B conformational landscape for this conserved active-site loop.

The work reported here validates local HDX-MS as a powerful method for elucidating detailed patterns of protein conformational dynamics and how they respond to binding of distinct ligands, complementing other methods such as time-resolved crystallography which can reveal altered enzyme dynamics in the presence vs. absence of small-molecule inhibitors ^25^. Future work can expand on this study by using HDX-MS to explore other allosteric ligands for PTP1B such as MSI-1436 ^13^, DPM-1001 ^26^, K197C covalent inhibitors ^11^, ABDF ^27^, and inhibitor 28p (initially designed for CD45 but found to have better affinity for PTP1B) ^28^. Investigating the impact of post-translational modifications (PTMs) and other binding proteins on PTP1B dynamics would also provide valuable insights into broader regulatory mechanisms. In addition, future studies may benefit from use of longer constructs of PTP1B that include more of the disordered C-terminus, which contains regulatory elements such as PTM sites and can harbor small-molecule allosteric inhibitors ^13^ but remains relatively poorly understood.

Finally, a potential avenue for future research involves using integrative computational methods to predict protein structural ensembles based on HDX patterns. Our preliminary analysis here using a pseudo-ensemble of many crystal structures (**Fig. 3**) hints at the potential of this approach. Moving forward, by combining HDX data with other types of computational modeling ^16^, it may be possible to develop more accurate and predictive models that incorporate dynamic information. This approach holds promise for advancing our understanding of protein dynamics and aiding in designing novel therapeutics targeting PTP1B and other proteins of biomedical interest.

## Methods

### Protein expression and purification

The protein purification method employed in this study was based on the purification protocol previously described ^29^, with slight modifications. We used the WT* construct of PTP1B, incorporating the C32S/C92V double mutation and including residues 1-321, for consistency with our previous crystallographic studies ^11^. This construct employs a pET24b vector containing a kanamycin resistance gene.

### Expression

To initiate protein expression, BL21 *E. coli* cells were transformed with the plasmid and cultured overnight at 37°C on LB agar plates supplemented with 35 µg/mL kanamycin. Subsequently, 5 mL starter cultures of LB medium containing 35 µg/mL kanamycin were inoculated with individual colonies and incubated overnight at 37°C with shaking at 170 rpm. These starter cultures were then used to inoculate larger 1 L cultures of LB medium supplemented with 35 µg/mL kanamycin. The cultures were grown at 37°C with shaking until the optical density at 600 nm reached approximately 0.6-0.8. Protein expression was induced by the addition of IPTG with a final concentration of 100 mM, followed by further incubation with shaking for either 4 hours at 37°C or overnight at 18°C. The resulting cell pellets were harvested by centrifugation and stored at −80°C in 50 mL conical tubes.

### Purification

The cell pellets were resuspended in lysis buffer (100 mM MES at pH 6.5, 1 mM EDTA, and 1 mM DTT) plus protease inhibitor and the solution was sonicated and centrifuged for 45 min at 20,000 RPM while kept at 4°C to obtain a clarified solution of cell lysate. This clarified sample was filtered (0.2 µm) and loaded onto a cation exchange chromatography SP FF 16/10 column (GE Healthcare Life Sciences) using the AKTA FPLC system. PTP1B eluted at approximately 200 mM NaCl in a multi-stage gradient of 0-1 M NaCl, with the gradient becoming steeper to facilitate elution of the target protein. Subsequently, size exclusion chromatography was performed using a Superdex 75 column (GE Healthcare Life Sciences) equilibrated with a crystallization buffer containing 10 mM Tris pH 7.5, 0.2 mM EDTA, 3 mM DTT, and 25 mM NaCl. The purity of the obtained PTP1B protein was assessed by analyzing samples on SDS-PAGE gels, demonstrating its high level of purity.

### Local HDX-MS experiments

#### Sample Handling

For each experiment, liquid handling was performed by the LEAP HDX platform. This robotic system precisely initiates and times labeling reactions (reaction volume of 50 µL), followed by rapid mixing with quench solution and dropping of temperature to 0-4°C. Once the sample was thoroughly mixed with quench, 100 µL of this quenched sample was then injected into the pepsin column.

#### Digestion Optimization

Digestion optimization experiments were carried out to determine the optimal quench conditions to be used to both halt hydrogen-deuterium exchange and prepare the protein for digestion over a pepsin column by inducing partial or extensive denaturing / unfolding. A series of quenching solutions consisting of 1.5% formic acid, 3.0% acetonitrile, and varying concentrations (0, 0.5, 1.0, 2.0, 4.0 M) of guanidinium hydrochloride (GuHCl) was mixed in a 1:1 ratio with the unlabeled protein. The deuteration buffer was identical to the protein buffer used in the non-deuterated experiments, except for the presence of deuterium in high (>99.5%) abundance. To perform local hydrogen-deuterium exchange (HDX) experiments, a Waters Enzymate BEH Pepsin Column was employed to generate peptides for subsequent analysis. Peptic peptides were eluted through a C18 analytical column (Hypersil Gold, 50 mm length × 1 mm diameter, 1.9 μm particle size, Thermo Fisher Scientific) into a Bruker maXis-II ESI-QqTOF high-resolution mass spectrometer. Peptide maps were generated for PTP1B for each of the varied GuHCl conditions, and the best coverage and highest resolution was seen with conditions 2.0 M and 4.0 M. A concentration of 3.0 M of GuHCl was chosen for future experiments.

#### HDX Labeling

Purified apo protein (PTP1B 1-321 WT*) and proteins with saturated active-site inhibitor (TCS401 [100 µM] in 2% DMSO) and saturated allosteric inhibitor (BB3 [100 µM] in 2% DMSO) were prepared as described in the previous section. All experiments were carried out in the crystallization buffer at 15°C in order to best match pH, salt, and reducing conditions that are present in comparable structural studies of PTP1B. The protein sample was first diluted to 20 µM in the H_2_O crystallization buffer + 2% DMSO + 100 µM of inhibitor or no inhibitor. The protein was then mixed with the labeling (D_2_O) buffer identical in chemical composition to the H_2_O crystallization buffer, including the inclusion of 2% DMSO and, in the case of the saturated ligand-bound experiments, 100 µM of inhibitor compound. This was done in order to keep the DMSO and ligand content the same from the initiation of label mixing until quench to avoid any momentary escape from equilibration of the bound protein:ligand fraction. The labeling reaction mixture consisted of 1 part protein (20 µM in H_2_O crystal buffer) and 9 parts D_2_O crystal buffer for a total D_2_O content of 90% for the duration of the reaction. After timepoints of 30 s, 100 s, 300 s, 1000 s, 3000 s, and 10,000 s, the reaction was quenched. To stop the HDX labeling process, cold quenching solution (1.5% formic acid, 3.0% acetonitrile, and 3 M GuHCl) was mixed in a 1:1 ratio with the labeled sample. 100 µL of this solution was injected by the LEAP HDX system to a pepsin column for digestion and further analysis. MS/MS fragmentation was used to confirm peptide identity along the sequence of PTP1B 1-321 WT* and these peptides and their retention times were used by the HDExaminer software to identify them automatically in the ‘on-exchange’ experiments.

In order to correct for back exchange in the regular and quantitative experiments, fully deuterated (FD) preparations were carried out on apo WT* PTP1B. The protein was incubated at room temperature (20°C) for 8 days in 94% D_2_O crystallization buffer. Duplicate (n=2) experiments were run for the 30 s, 100 s, and 10,000 s time points, while single experiments were run for the intermediate time points of 300 s, 1000 s, and 3000 s. Triplicate (n=3) experiments were run for the quantitative HDX experiments for comparison with crystallographic metrics.

### HDX-MS data analysis

#### Data Preprocessing

Before performing the HDX analysis, the raw mass spectrometry data files (.d format) were processed using Compass Data Analysis 5.3 and Biotools 3.2 software to convert the data into a suitable format (.csv). The preprocessed data files were imported into version 3.3 of the HDExaminer software from Sierra Analytics. For accurate analysis, the imported data files were aligned based on the peptide identification information, retention time, and m/z values. This step ensures that the corresponding peptide measurements from different time points are properly aligned for further analysis. The imported data were matched with the peptide sequences derived from the pepsin-based protein digestion within HDExaminer.

#### Peptide-Level Exchange Rate Analysis

The deuteration level of each peptide was determined by comparing the centroid mass of the deuterated peptide ion with that of the corresponding non-deuterated peptide ion at each time point. This calculation provides the deuteration percentage for each peptide at different time intervals. The HDExaminer software employs various algorithms to calculate the exchange rates of individual peptides. These algorithms utilize mathematical models, such as exponential fitting, to estimate the exchange kinetics and determine the exchange rates of the identified peptides. Relative deuterium uptake is expressed as the peptide mass increase divided by the number of peptide backbone amides. When the second residue of the peptide is not proline, the number of peptide backbone amides was decreased by one to account for rapid back exchange by the amide adjacent to the N-terminal residue.

#### Visualization and Data Interpretation

The exchange rate profiles at the peptide and residue levels were computed in HDExaminer. The software provides graphical representations, such as heat maps and exchange rate plots, to facilitate the interpretation of the data. The exchange rate profiles can be further analyzed to identify regions of the protein exhibiting differential exchange behavior under different experimental conditions. To qualitatively visualize exchange values mapped to 3D protein structures, we obtained estimates of deconvoluted and smoothed residue-level interpolation of the peptide results from HDExaminer and plotted values along color spectra for %deuteration (**Fig. 2B**) and Δ%deuteration (**Fig. 5**) that were then mapped onto the solved structures for the apo and liganded conditions, respectively.

#### Data Availability

Data used in this paper including peptide raw centroids, quantitative HDX data, and crystallographic Cα RMSFs are available as supplemental information in **Data S1**.

### PTP1B crystal structures pseudo-ensemble

Crystal structures of PTP1B were obtained from the Protein Data Bank (PDB) by searching for all structures with 95% or greater sequence identity to PDB ID 1SUG, excluding structures from PanDDA^21^. The final list of structures consisted of the following 199 PDB IDs: 1A5Y, 1AAX, 1BZC, 1BZH, 1BZJ, 1C83, 1C84, 1C85, 1C86, 1C87, 1C88, 1ECV, 1EEN, 1EEO, 1G1F, 1G1G, 1G1H, 1G7F, 1G7G, 1GFY, 1I57, 1JF7, 1KAK, 1KAV, 1L8G, 1LQF, 1NL9, 1NNY, 1NO6, 1NWE, 1NWL, 1NZ7, 1OEM, 1OEO, 1OES, 1OET, 1OEU, 1OEV, 1ONY, 1ONZ, 1PA1, 1PH0, 1PTT, 1PTU, 1PTV, 1PTY, 1PXH, 1PYN, 1Q1M, 1Q6J, 1Q6M, 1Q6N, 1Q6P, 1Q6S, 1Q6T, 1QXK, 1SUG, 1T48, 1T49, 1T4J, 1WAX, 1XBO, 2AZR, 2B07, 2B4S, 2BGD, 2BGE, 2CM2, 2CM3, 2CM7, 2CM8, 2CMA, 2CMB, 2CMC, 2CNE, 2CNF, 2CNG, 2CNH, 2CNI, 2F6F, 2F6T, 2F6V, 2F6W, 2F6Y, 2F6Z, 2F70, 2F71, 2FJM, 2FJN, 2H4G, 2H4K, 2HB1, 2HNP, 2HNQ, 2NT7, 2NTA, 2QBP, 2QBQ, 2QBR, 2QBS, 2VEU, 2VEV, 2VEW, 2VEX, 2VEY, 2ZMM, 2ZN7, 3A5J, 3A5K, 3CWE, 3D9C, 3EAX, 3EB1, 3EU0, 3I7Z, 3I80, 3QKP, 3QKQ, 3SME, 3ZMP, 3ZMQ, 3ZV2, 4BJO, 4I8N, 4QAH, 4QAP, 4QBE, 4QBW, 4Y14, 4ZRT, 5K9V, 5K9W, 5KA0, 5KA1, 5KA2, 5KA3, 5KA4, 5KA7, 5KA8, 5KA9, 5KAA, 5KAB, 5KAC, 5KAD, 5T19, 6B8E, 6B8T, 6B8X, 6B8Z, 6B90, 6B95, 6BAI, 6CWU, 6CWV, 6NTP, 6OL4, 6OLQ, 6OLV, 6OMY, 6PFW, 6PG0, 6PGT, 6PHA, 6PHS, 6PM8, 6W30, 6XE8, 6XEA, 6XED, 6XEE, 6XEF, 6XEG, 7KEN, 7KEY, 7KLX, 7L0C, 7L0H, 7LFO, 7MKZ, 7MM1, 7MN7, 7MN9, 7MNA, 7MNB, 7MNC, 7MND, 7MNE, 7MNF, 7MOU, 7MOV, 7MOW, 7RIN, 7S4F, 8DU7, 8G65, 8G67, 8G68, 8G69, 8G6A. The Python package ProDy ^30^ was used to align the structures, construct the pseudo-ensemble, and calculate Cα RMSF values.

## Acknowledgments

DAK is supported by NIH R35GM133769.

We thank Simeng (Selina) Sun for calculating the Cα RMSF values, and Shawn Michael Costello for helpful comments on the manuscript.

