## Supplemental Figures for "Native dynamics and allosteric responses in PTP1B probed by high-resolution HDX-MS"

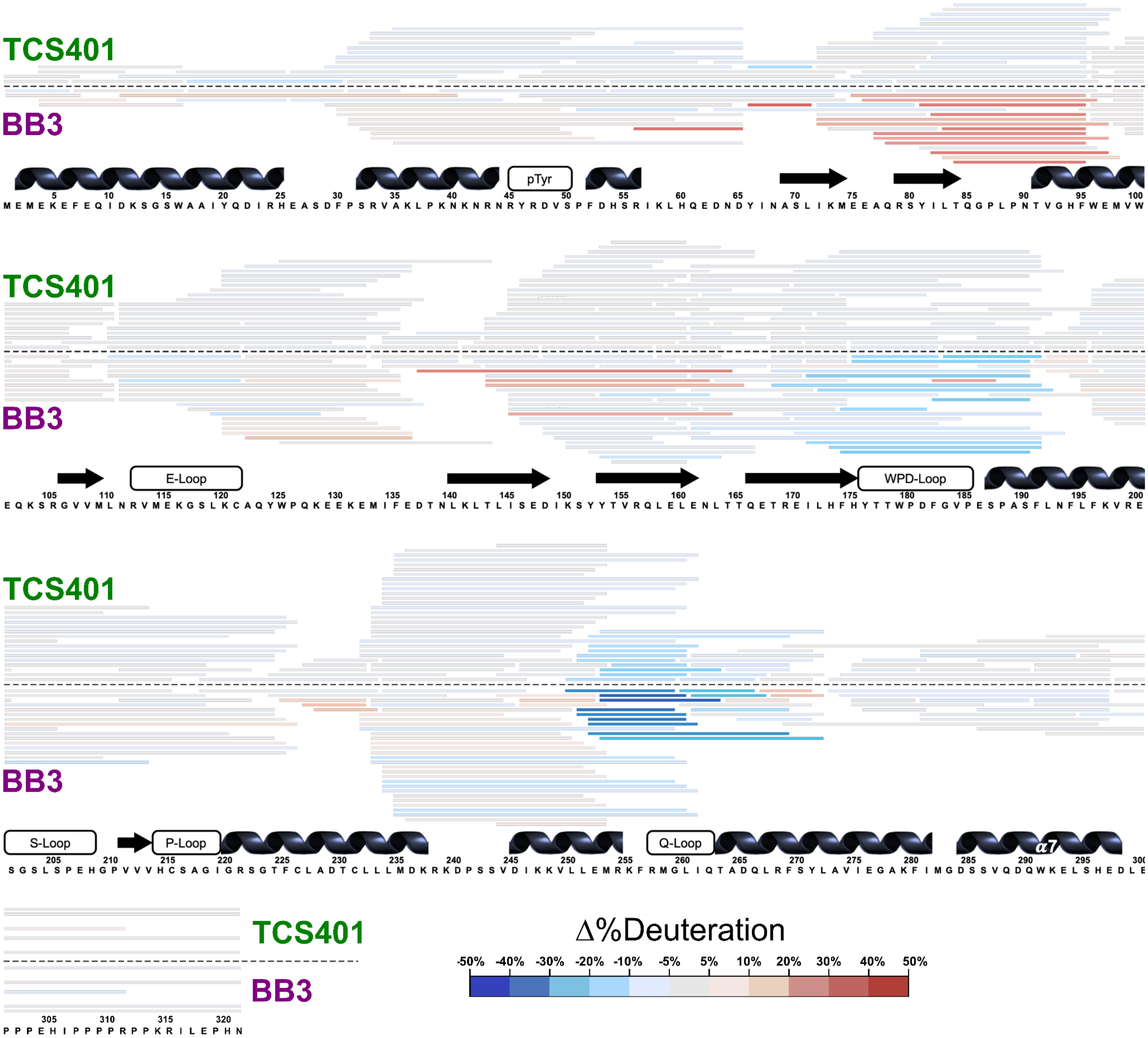


**Figure S1: Difference HDX map for TCS401 minus apo PTP1B.**
Peptide maps for the TCS401 and BB3 conditions, with peptides colored based on the difference in HDX rate between liganded minus apo conditions.

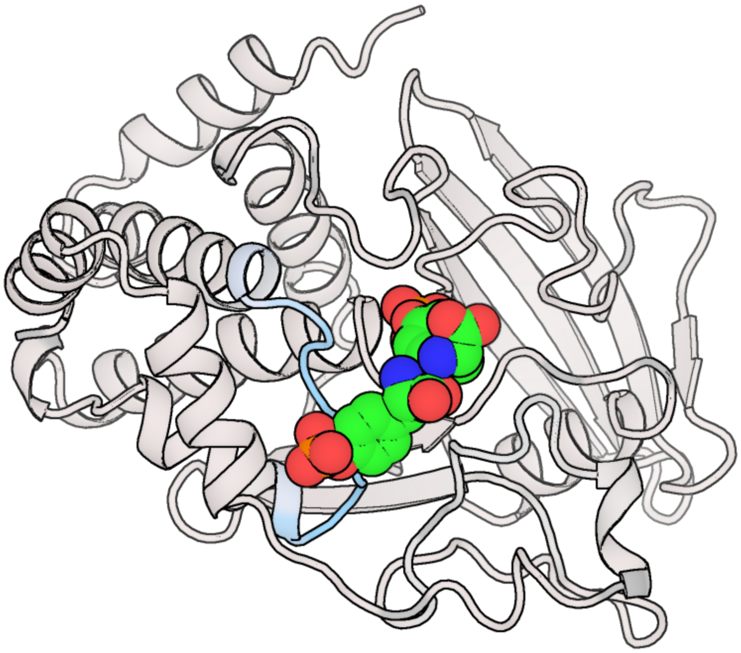

**Figure S2: Structural view of TCS401 HDX differences in context of two pTyr sites.**
Same as main **Fig. 5A** but with TCS401 omitted and the two pTyr residues from PDB ID 1PTY overlaid (green).

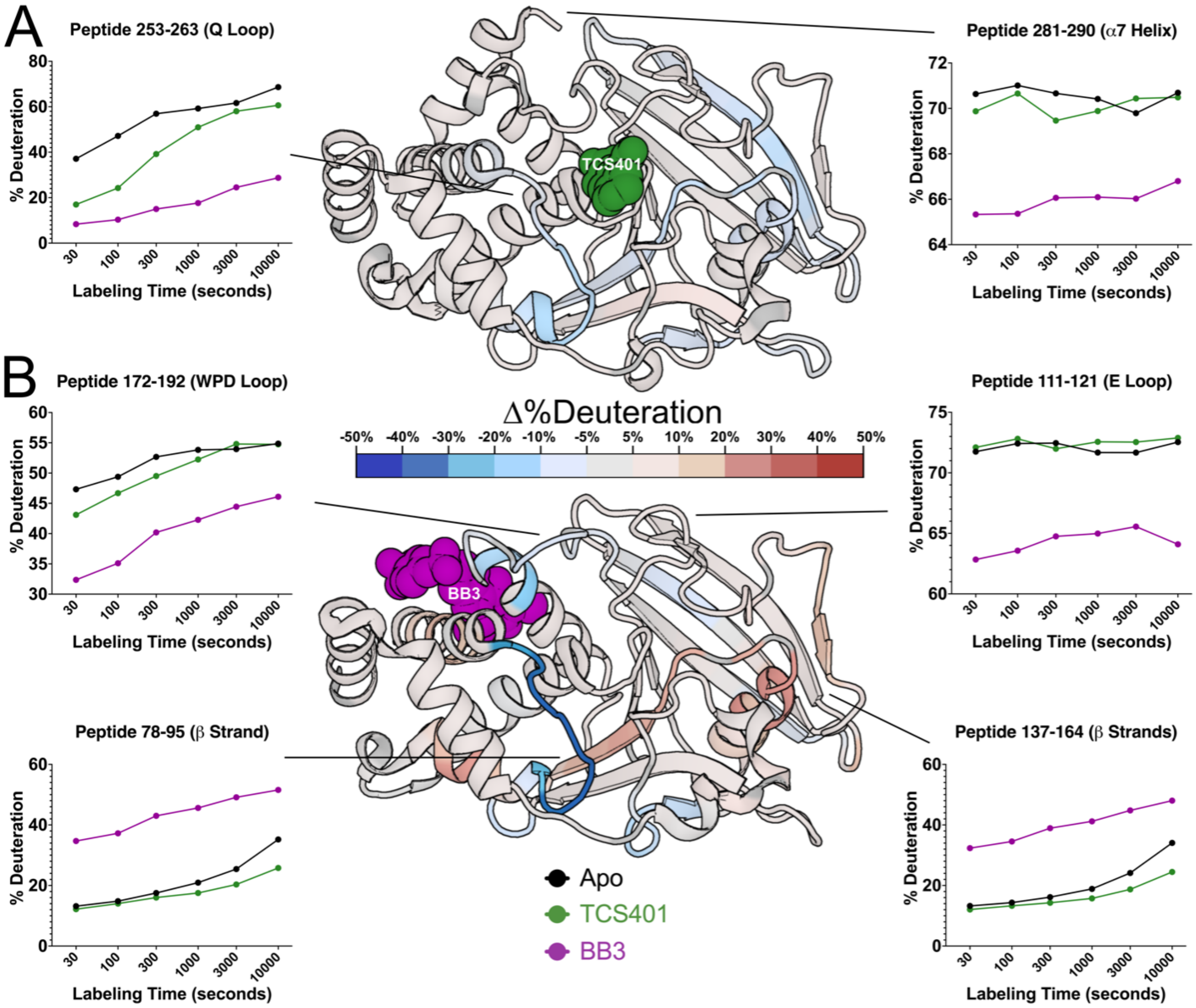


**Figure S3: Effects of active-site inhibitor (TCS401) and allosteric-site inhibitor (BB3) after extended labeling time.**

Same as **Fig. 5** but with coloring of residues at 10,000 seconds of labeling time.

**Data S1: Fractional deuterium uptake for all three conditions.**

Spreadsheet containing HDX and pseudo-ensemble data.

Tab 1: Peptide raw centroids for all three protein conditions across all time points.

Tab 2: Quantitative HDX data for apo PTP1B at 30 seconds of labeling time.

Tab 3: Cα RMSFs from pseudo-ensemble of PTP1B structures.
